# Interferon induction and not replication interference mainly determines anti-influenza virus activity of defective interfering particles

**DOI:** 10.1101/2021.09.20.461172

**Authors:** Prerna Arora, Najat Bdeir, Sabine Gärtner, Stefanie Reiter, Lars Pelz, Ulrike Felgenhauer, Udo Reichl, Stephan Ludwig, Friedemann Weber, Markus Hoffmann, Michael Winkler, Stefan Pöhlmann

## Abstract

Defective interfering (DI) RNAs arise during influenza virus replication, can be packaged into particles (DIPs) and suppress spread of wildtype (WT) virus. However, the molecular signatures of DI RNAs and the mechanism underlying antiviral activity are incompletely understood. Here, we show that any central deletion is sufficient to convert a viral RNA into a DI RNA and that antiviral activity of DIPs is inversely correlated with DI RNA length when induction of the interferon (IFN) system is disfavored. When induction of the IFN system was allowed, it was found to be the major contributor to DIP antiviral activity. Finally, while both DIPs and influenza virus triggered expression of IFN-stimulated genes (ISG) only virus stimulated robust expression of IFN. These results suggest a key role of innate immune activation in DIP antiviral activity and point towards previously unappreciated differences in DIP- and influenza virus-mediated activation of the effector functions of the IFN system.

**Importance:** Defective interfering (DI) RNAs naturally arise during RNA virus infection. They can be packaged into defective interfering particles (DIPs) and exert antiviral activity by suppressing viral genome replication and inducing the interferon (IFN) system. However, inhibition of influenza virus infection by DI RNAs has been incompletely understood. Here, we show that induction of the IFN system and not suppression of genome replication is the major determinant of DIP antiviral activity. Moreover, we demonstrate that DIPs induce IFN-stimulated genes (ISG) but not IFN with high efficiency. Our results reveal unexpected major differences in influenza virus and DIP activation of the IFN system, a key barrier against viral infection, and provide insights into how to design DIPs for antiviral therapy.

## Introduction

The annually recurring influenza epidemics are a major source of global morbidity and mortality and intermittent pandemics can have even more severe consequences (1). Influenza therapy and vaccination are available but suffer from serious shortcomings (1). The success of influenza therapy with currently licensed drugs, which target the viral proteins neuraminidase (NA), matrix protein 2 (M2) or polymerase acidic protein (PA), can be compromised by resistance development (2). Moreover, vaccines against epidemic influenza need to be annually adjusted to the viruses expected to circulate during the next influenza season and offer little or no protection against emerging pandemic viruses (1). Thus, the identification of novel targets and strategies for antiviral intervention is an important task.

Influenza viruses contain a segmented, negative sense RNA genome. The genomic segments are replicated by the viral polymerase, which consists of the subunits polymerase basic proteins 1 (PB1) and 2 (PB2) as well as PA (3). The error rate of the viral polymerase is high and can result in the synthesis of genomic segments that harbor deletions (4-10). These defective segments may interfere with the amplification of wt segments and are thus termed defective interfering (DI) RNAs (4-6, 8-10). Packaging of DI RNAs into viral particles results in the formation of DI particles (DIPs), which suppress wt influenza virus spread (4, 5). It has been proposed that DIPs suppress influenza virus infection by interfering with genome replication (a process subsequently termed replication interference) and by inducing an interferon (IFN) response (4, 5, 11-17). However, this concept has not been systematically investigated and the relative contribution of replication interference and IFN induction to DIP antiviral activity is unknown.

We recently developed a cell culture system that allows production of genetically defined DIPs based on reverse genetics and a cell line complementing defects in influenza A virus (IAV) genomic segment 1 (18). Here, we used this system as well as a mini-replicon assay (19) to analyze the contribution of replication interference and IFN induction to the antiviral activity of DIPs. We report that in the mini-replicon assay any central deletion in segment 1, 2 or 3 converts these segments into DI RNAs, which suppress replication of diverse target segments. Inhibitory activity of these DI RNAs was inversely correlated with segment length and a similar correlation was seen in the context of DIP and IAV infection under conditions which disfavored IAV inhibition by DIP-dependent induction of the IFN system. If induction of the IFN system was allowed before IAV infection, it largely accounted for DIP antiviral activity. Finally, DIPs robustly induced IFN-stimulated gene (ISG) but not IFN expression, indicating that IAV and DIPs may differ in the activation of the effector functions of the IFN system. Our results suggest that although interference with genome replication contributes to DIP antiviral activity, the induction of the IFN system is the major determinant of suppression of virus infection by DIPs.

## Results

### DI-244 inhibits segment replication in a mini-replicon assay and inhibition is independent of the truncated PB2 open reading frame

We first investigated whether a previously described IAV mini-replicon assay (19) is suitable to detect inhibition of IAV genome replication by a prototypic segment 1-derived DI RNA, DI-244 (20). This assay is based on a firefly luciferase open reading frame flanked by the 5’ and 3’ ends of IAV segment 8, which is amplified in cells upon coexpression of the constituents of the viral polymerase complex PB1, PB2, and PA, and the viral nucleoprotein (NP) (19). Transfection of 293T cells with plasmids encoding the mini-genome reporter segment and the IAV proteins mentioned above resulted in luciferase activities in cell lysates that were approximately 1,000-fold higher than those measured in cells transfected with the reporter alone or transfected with the full set of plasmids except the PB2 encoding plasmid (Figure 1A). Moreover, cotransfection of two different amounts of DI-244 encoding plasmid resulted in a concentration dependent decrease in luciferase activity, indicating that DI-244 inhibited replication of the reporter segment (Figure 1A). This inhibitory activity was also observed when the PB2 start codon in DI-244 and two subsequent ATGs (positions 11 and 28) were mutated (Figure 1B). In contrast, transfection of expression plasmid pCAGGS containing the truncated PB2 ORF of DI-244 or empty pCAGGS did not reduce luciferase signals (Figure 1B). These results indicate that inhibition of segment replication by DI-244 can be visualized in the mini-replicon assay and does not require expression of truncated PB2.

**FIG 1.**
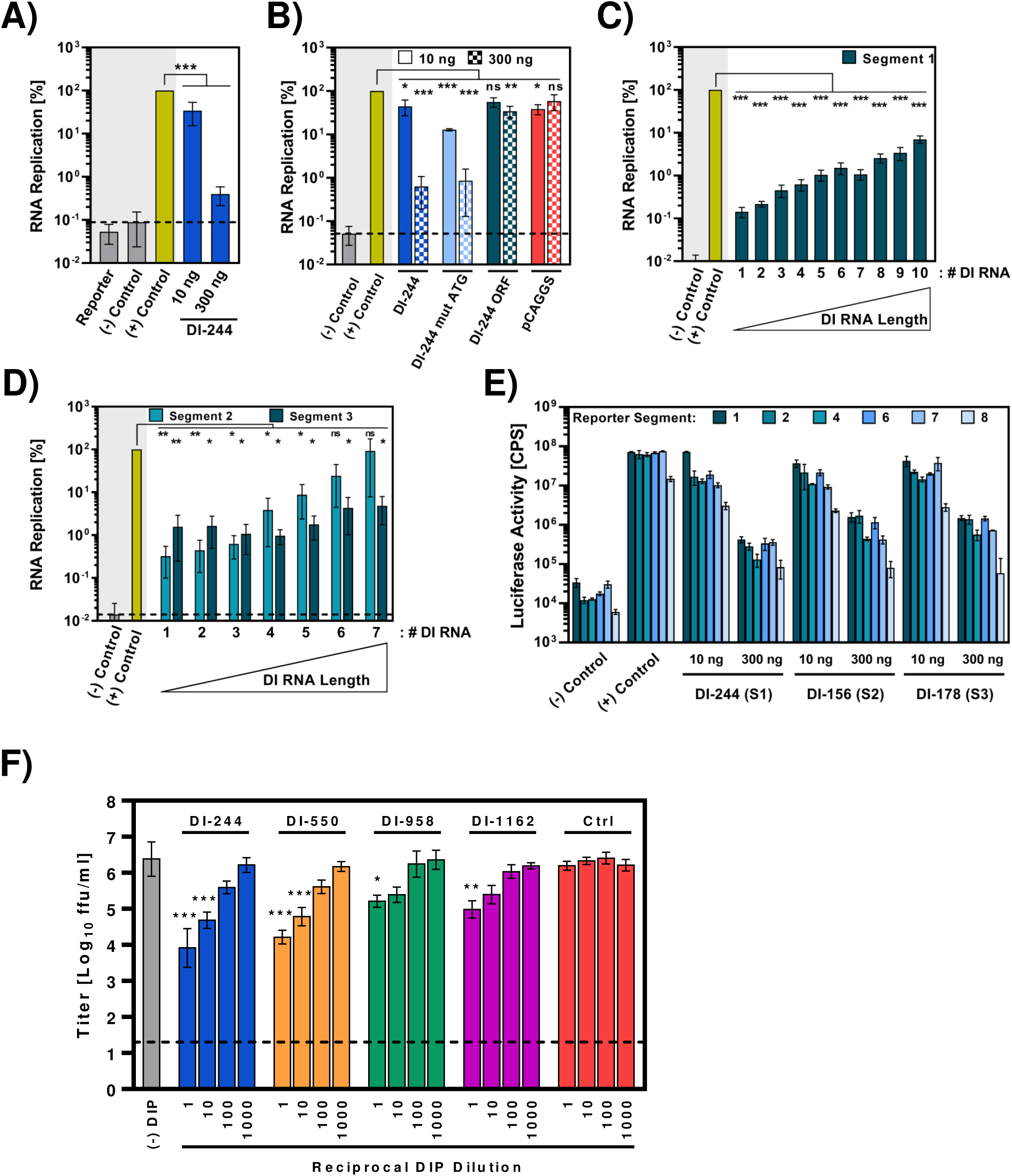
Antiviral activity of DI RNAs inversely correlates with DI RNA length (A) DI-244 inhibits genome replication in the mini-replicon assay. 293T cells were transfected with plasmids encoding the viral polymerase proteins, NP, a segment 8-based luciferase reporter (mini-replicon system) and either empty plasmid or plasmid for expression of DI-244 mRNA (10 and 300 ng) and vRNA. Removing the plasmid encoding PB2 from the transfection mix served as negative control. Cotransfection of all support plasmids and empty plasmid instead of DI-244 encoding plasmid served as positive control. The average of five independent experiments is shown, for which the positive control was set as 100%. Error bars indicate standard error of the mean (SEM). (B) The truncated open reading frame of DI-244 does not contribute to inhibition of genome replication in the mini-replicon assay. The experiment was carried out as described for panel A but the cells were cotransfected with a plasmid for expression of DI-244 mRNA and vRNA with or without the first three ATGs of the PB2 ORF being intact (DI-244, DI-244 mut ATG), a plasmid for expression of DI-244 mRNA (DI-244 ORF) or empty plasmid pCAGGS. The average of three independent experiments is shown, for which the positive control was set as 100%. Error bars indicate SEM. (C) The inhibitory activity of segment 1-derived DI RNAs in the mini-replicon assays is inversely correlated with DI RNA length. The experiment was carried out as described for panel A but 300 ng of plasmids harboring the indicated segment 1-derived DI RNAs were cotransfected. The DI RNAs tested were numbered as shown in table S1. The average of five independent experiments is shown, for which the positive control was set as 100%. Error bars indicate SEM. (D) The inhibitory activity of segment 2- and 3-derived DI RNAs in the mini-replicon assays is inversely correlated with DI RNA length. The experiment was conducted as described for panel A but 300 ng of plasmids harboring the indicated segment 2 and 3-derived DI-RNAs were cotransfected. The DI RNAs tested were numbered as shown in table S1. The average of three independent experiments is shown, for which the positive control was set as 100%. Error bars indicate SEM. (E) The inhibitory activity of DI RNAs in the mini-replicon assays is independent from the origin of the reporter segment. The experiment was carried out as described for panel A but the indicated reporter segments and segment 1, 2 and 3-derived DI RNAs were used. The results of a single representative experiment are shown and were confirmed in an independent experiment. Error bars indicate standard deviation (SD). (F) Antiviral activity of segment 1-derived DIPs is inversely correlated with DI RNA lengths in the presence of trypsin. MDCK cells were coinfected with the indicated DIPs (MOI 1) and A/PR/8/34 (MOI 0.001) in the presence of trypsin, washed, and cultured in medium with trypsin. DIP-negative supernatants served as controls. At 72 h post infection, viral titers in culture supernatants were determined by focus formation assay. The average of four independent experiments is shown; error bars indicate SEM. In panels A-D statistical significance of differences between values measured for cells cotransfected with support plasmids and either empty plasmid (+ control) or DI RNA encoding plasmid was determined using one-way ANOVA with Sidak’s (panel A) or with Dunnett’s posttest (panel B-D). In panel F statistical significance of differences between values measured for cells infected with virus and cells infected with virus in the presence of DIPs was determined using one-way ANOVA with Dunnett’s posttest. *, p ≤ 0.05; **, p ≤ 0.01; ***, p ≤ 0.001

### Inhibitory activity of segment 1, 2 and 3-derived DI RNAs is inversely correlated with RNA length and is independent of the target segment

It is believed that the short length of DI-244, as compared to wt segment 1, results in faster amplification of DI-244 and ultimately in suppression of amplification of the wt segment (4, 5). If correct, one would assume that DI RNA length is a major determinant of antiviral activity. We explored this possibility by investigating the capacity of a set of ten segment 1-derived RNAs with nested central deletions to inhibit segment amplification in the mini-replicon assay. All RNAs tested exerted inhibitory activity and an inverse correlation between RNA length and inhibitory activity was observed (Figure 1C, Table S1). When the plasmids encoding the DI RNAs with the smallest and largest deletion were normalized for copy numbers instead of weight little effect on inhibitory activity was observed. Thus, the inverse correlation between DI RNA length and inhibitory activity was not due to differences in plasmid copy numbers transfected (not shown). Finally, further shortening of DI-244 did not augment inhibitory activity (not shown), suggesting that DI-244 length may be optimal for inhibition of wt segment replication. In sum, our results show that the ability of segment 1-derived DI RNAs to block replication of a wt segment is dependent on the DI RNA length.

We next explored whether the inverse correlation between length and inhibitory activity is also observed for segment 2- and 3-derived DI RNAs. For this, we introduced central, nested deletions in segment 2 and 3 and investigated inhibitory activity in the mini-replicon system. As for segment 1-derived RNAs, all segment 2- and 3-based RNAs with deletions exerted inhibitory activity and inhibition inversely correlated with RNA length, although this correlation was more pronounced for segment 2 as compared to segment 3 (Figure 1D, Table S1).

Next, we examined whether the segment 1-, 2- and 3-derived DI RNAs with the largest deletion (constructs DI-244 (segment 1, S1), DI-156 (segment 2, S2), DI-178 (segment 3, S3), Table S1) were able to efficiently suppress replication of different IAV segments or were mainly active against segment 8, which was so far employed in the mini-replicon assay. For this, we added the 5’ and 3’ ends of segments 1, 2, 4, 6, and 7 to the firefly luciferase sequence and tested the amplification of these reporter segments in the mini-replicon assay. In the absence of DI RNAs, all segments were efficiently amplified, as demonstrated by high luciferase activity in lysates of cells coexpressing PB2, PB1, PA and NP (Figure 1E). Cotransfection of two different amounts of segment 1-, 2- or 3-derived DI RNAs reduced replication of all reporter segments efficiently and in a concentration dependent manner (Figure 1E). Thus, in the mini-replicon assay, introduction of a deletion into an IAV genomic segment is sufficient to convert it into a DI RNA and length and inhibitory activity of these DI RNAs are inversely correlated.

### Inverse correlation between anti-IAV activity of DIPs and DI RNA length

We recently reported a cell culture system for production of DIPs in the absence of helper virus, which relies on IAV reverse genetics and DIP producer cell lines stably expressing the PB2 protein (18). We employed this system to generate DIPs with nested deletions in segment 1 and assessed their ability to inhibit infection of MDCK cells with A/PR/8/34 (PR8). We found that DI-244, which contains the smallest DI RNA, inhibited PR8 infection with the highest efficiency and that inhibitory activity of DIPs decreased as DI RNA length increased (Figure 1F). Thus, an inverse correlation between DI RNA length and inhibitory activity observed in the mini-replicon assay could be confirmed in the context of DIPs, at least under the conditions tested.

### Preincubation of target cells with DI-244 increases antiviral activity

It has been reported that DIPs can block viral infection by stimulating the IFN system (11, 12). Therefore, we sought to clarify whether induction of the IFN system could contribute to DI-244 antiviral activity in MDCK cells. Trypsin is used for A/PR8/34 activation but can inactivate IFNα (Figure 2A) (21) and can thus cofound analyses of IAV inhibition by the IFN system. Therefore, we switched to A/WSN/33 (WSN) as challenge virus and WSN-derived DIPs, since WSN can replicate trypsin-independently in cell cultures containing fetal bovine serum (FBS) (22). To obtain first insights into a potential role of the IFN system in DIP antiviral activity, we reasoned that if induction of the IFN system was a major determinant of DIP antiviral activity, then time-of-DIP addition to target cells should have a major impact on the efficiency of IAV inhibition by DIPs. Thus, addition of DIPs and virus to target cells at the same time should preclude the establishment of a robust DIP-induced antiviral state prior to IAV infection. In contrast, addition of DIPs at 24 h before virus should allow for establishment of such an antiviral state and might thereby boost DIP antiviral activity. Preincubation of target MDCK cells with DI-244 for 24 h indeed increased DIP antiviral activity as compared to simultaneous addition of DI-244 and IAV, especially when high doses of DI-244 were analyzed (Figure 2B, left panel). Unexpectedly, similar results were obtained in the presence of trypsin (Figure 2B, right panel), indicating that the enhanced antiviral activity of DI-244 upon 24 h preincubation with target cells was likely not due to induction of IFNα or another trypsin-sensitive antiviral host cell protein.

**FIG 2.**
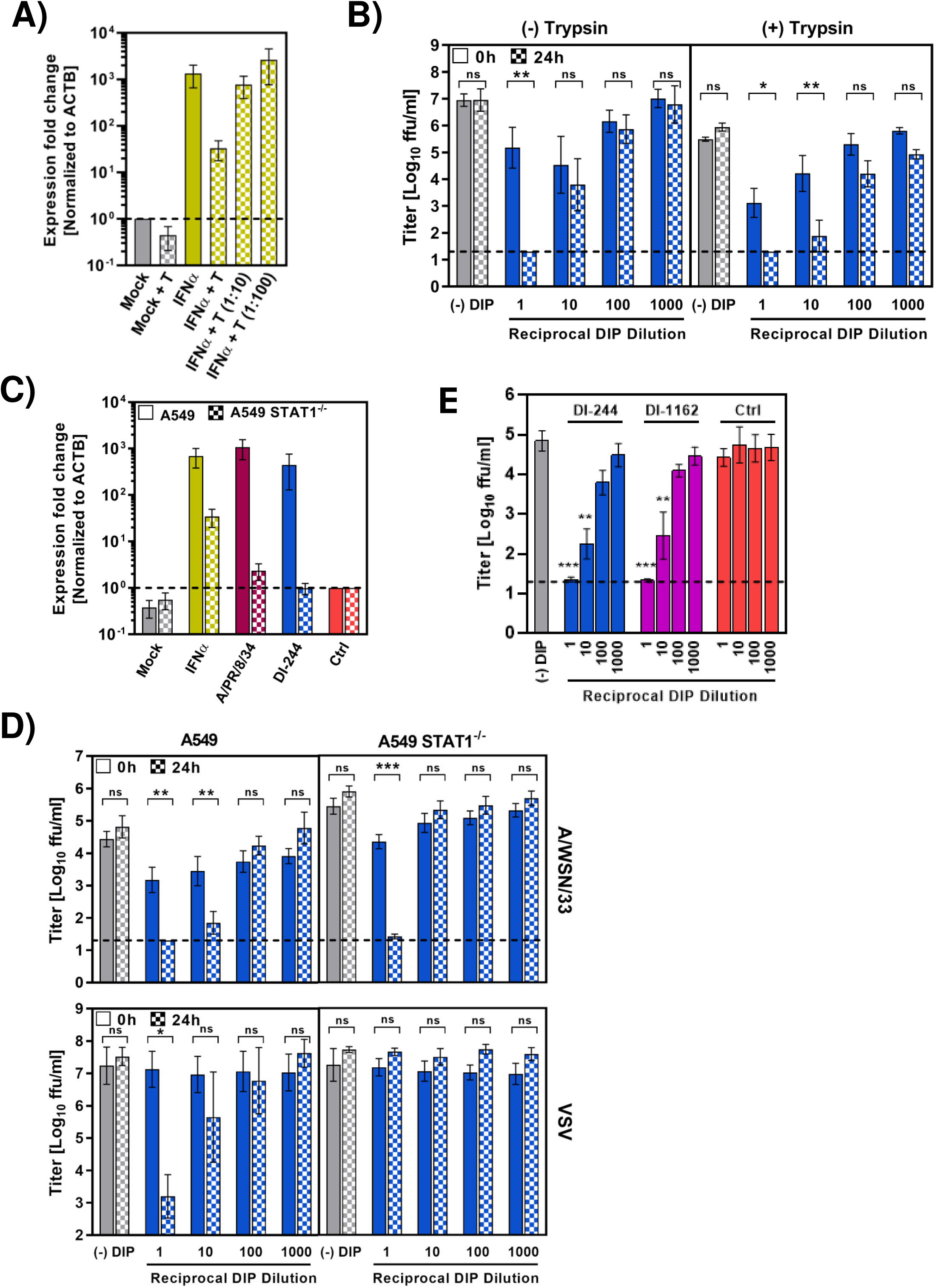
Induction of the IFN system is a major contributor to DIP antiviral activity (A) Trypsin inactivates IFNα. A549 wt cells were exposed to recombinant IFNα (100 U/ml) in the presence and absence of serially diluted trypsin (T). Undiluted trypsin (IFNα + T) was added at a concentration of 0.5 µg/ml. After 24 h, cells were harvested, RNA isolated and *MX1* expression analyzed by quantitative RT-PCR. *MX1* transcripts levels were normalized against *ACTB (β-actin)* transcript levels. The average of three independent experiments is shown. Error bars indicate SEM. (B) Pre-exposure of target cells to DIPs boosts DIP antiviral activity independent of trypsin. Left panel, - Trypsin condition: MDCK cells were either coinfected with DI-244 (MOI 10) and A/WSN/33 (MOI 0.1) in the absence of trypsin or DI-244 was added to cells at 24 h before virus. Cells were washed 1 h after addition of virus and maintained in growth medium without trypsin. At 72 h post infection, viral titers in culture supernatants were determined by focus formation assay. Right panel, + trypsin condition: The experiment was carried out as described for the left panel, but A/WSN/33-derived DIPs (MOI 1) and A/WSN/33 (MOI 0.001) were used and maintained in infection medium supplemented with trypsin. The average of three independent experiments is shown in both panels; error bars indicate SEM. (C) STAT1 is required for *MX1* induction by IAV and DIP. A549 cells and A549 STAT1^-/-^ cells were exposed to IFNα (100 U/ml), A/PR/8/34 or DI-244 (all MOI 1, in the presence of trypsin) for 1 h, washed, incubated for 24 h in the absence of trypsin and *MX1* mRNA expression quantified using qRT-PCR. The average of five independent experiments is shown. Error bars indicate SEM. (D) Anti-IAV activity of DI-244 is partially and anti-VSV activity of DIP is fully dependent on STAT1. Antiviral activity of DI-244 was analyzed as described for the left panel of figure 2B but A549 wt and A549 STAT1^-/-^ cells were used. At 96 h post infection, viral titers in culture supernatants were determined by focus formation assay. The average of six (A/WSN/33) and three independent experiments (VSV) is shown. Error bars indicate SEM. (E) DI RNA length does not modulate DIP antiviral activity in the context of a functional IFN system. Antiviral activity of the indicated DIPs was analyzed as described for panel D adding DIPs 24 h before virus. The average of five independent experiments is shown. Error bars indicate SEM. In panels B and D statistical significance of differences between values measured for cells inoculated with DIPs at 24 h before IAV infection and cells to which IAV and DIPs were added at the same time was determined using two-way ANOVA with Sidak’s posttest. In panel E statistical significance of differences in viral titers obtained on cells treated with different concentrations of DIPs or without (-) DIPs was determined using one-way ANOVA with Dunnett’s posttest. *, p ≤ 0.05; **, p ≤ 0.01; ***, p ≤ 0.001

### DI-244 induces anti-IAV activity in A549 cells in a STAT1-independent fashion

In order to more directly assess the contribution of the IFN system to DI-244 antiviral activity, we employed A549 wt cells and A549 cells which lack STAT1 (signal transducer and activator of transcription 1, STAT1^-/-^) and are thus defective in IFN-induced signaling. Confirmatory experiments revealed that IFNα, IAV and DI-244 strongly upregulated *MX1* gene expression in A549 wt but not *STAT1*^-/-^ cells, in keeping with a defective JAK/STAT signaling pathway (Figure 2C). Addition of undiluted and 1:10 diluted DI-244 to A549 cells at 24 h before infection with WSN resulted in 100 -fold higher antiviral activity as compared to DI-244 added at the same time as virus (Figure 2D), confirming and extending the data obtained with MDCK cells.

Unexpectedly, addition of undiluted DIP to A549 STAT1^-/-^ cells still resulted in high antiviral activity (Figure 2D), although 10-fold diluted DI-244 showed markedly reduced antiviral activity in STAT1^-/-^ cells as compared to wt cells. In contrast, inhibition of vesicular stomatitis virus (VSV) infection by DI-244 was completely dependent on STAT1, independent of the DIP dose used (Figure 2D). Finally, we asked whether the antiviral activity of DIPs still depended on the DI RNA length if DIPs were added to cells before virus. In contrast to what was observed with MDCK cells in the presence of trypsin, all DIPs with nested deletions in segment 1 inhibited WSN infection of A549 wt cells with similar efficiency (Figure 2E and data not shown), indicating that the contribution of replication interference to DIP antiviral activity was minor or absent under those conditions. Collectively, our findings indicate that DIPs can induce robust, partially STAT1-independent anti-IAV activity that is not determined by DI RNA length and markedly more potent than DIP-mediated inhibition of IAV genome replication.

### DI-244 induces robust expression of ISGs but not IFN

In order to understand how DIPs activate the IFN system, we compared DIP- and IAV-mediated stimulation of IFN expression. For this, an IFN bioassay was employed that was based on VSV, a highly IFN-sensitive virus (23). A549 or A549 STAT1^-/-^ effector cells were incubated with IAV, VSV or DI-244 for 24 h, the supernatants collected and heat and acid treated to inactivate viral particles but not IFN, which is known to display a certain heat and acid stability. Subsequently, the supernatants were added to sentinel cells (A549) for 24 h followed by inoculation of the sentinel cells with a single-cycle reporter VSV replicon and quantification of infection. For standardization, A549 cells were incubated with recombinant IFNα, infected with the single-cycle VSV and infection efficiency quantified. Supernatants from IAV exposed A549 wt cells but not A549 STAT1^-/-^ cells potently inhibited subsequent VSV infection (Figure 3A), indicating that IAV induced production of IFN in a STAT1-dependent fashion, as expected. Similar findings were made with supernatants from VSV exposed cells but antiviral activity was independent of STAT1 expression (Figure 3A), again in agreement with published data (24). Finally, and unexpectedly, supernatants from A549 wt cells exposed to DI-244 were not inhibitory and the same finding was made for supernatant from DI-244 treated A549 STAT1^-/-^ cells, indicating that IFN induction by DI-244 was low or absent (Figure 3A).

**FIG 3.**
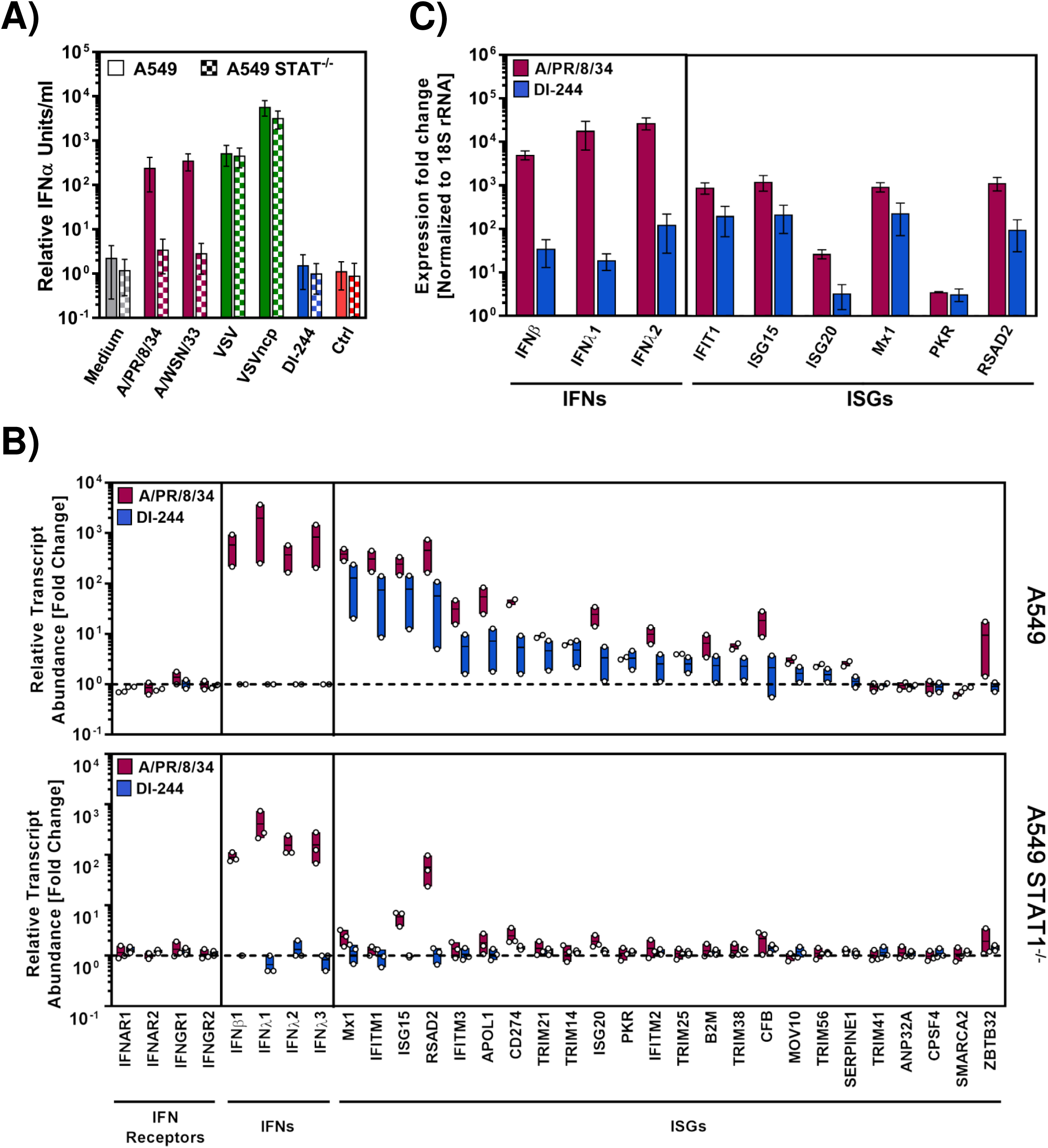
DI-244 robustly induces ISG but not IFN expression (A) DI-244 does not induce IFN expression as determined in a VSV-replicon-based bioassay. A549 and A549 STAT1^-/-^ effector cells were exposed to IAV, VSV or DI-244 and supernatants collected, heat inactivated, acid treated and added onto A549 sentinel cells followed by infection with VSV. For calibration, A549 cells were incubated with recombinant IFNα, VSV infected and infection efficiency was quantified. The average of three independent experiments is shown. Error bars indicate SEM. (B) DI-244 induces robust ISG but not IFN expression as determined by RNAseq. A549 cells (top panel) and A549 STAT1^-/-^ cells (bottom panel) were incubated with IAV (A/PR/8/34), DI-244 and control supernatants at an MOI of 1 in the absence of trypsin and subjected to RNAseq analysis. Expression of selected ISGs is shown. The average of two independent experiments (A549) and three experiments (A549 STAT1^-/-^) is presented. Error bars indicate SEM and black lines within the floating bars indicate the mean. (C) DI-244 induces robust ISG but not IFN expression as determined by qRT-PCR analysis. The A549 wt cells described in panel B were subjected to qRT-PCR analysis of ISG expression. The average of three independent experiments is shown.

The ability of DI-244 to inhibit IAV and VSV infection without inducing IFN, as determined in the bioassay, posed the question how DI-244 alters gene expression in target cells to block infection. To address this question, A549 cells and A549 STAT1^-/-^ were either incubated with control supernatants or supernatants containing DI-244 or IAV and subjected to RNAseq analysis. PR8 was employed for these studies, in order to limit viral replication to a single cycle (since no trypsin was present after virus inoculation) (25). Neither PR8 nor DI-244 induced the expression of IFN receptors (Figure 3B). In contrast, PR8 but not DI-244 induced expression of IFNβ and IFNλ (Figure 3B), in agreement with our results obtained in the bioassay. Despite the differential upregulation of IFNs by PR8 and DI-244 both induced the robust expression of antiviral ISGs, including MX1, IFITM1 and ISG15 in A549 wt cells, although induction by PR8 was more efficient than that observed for DI-244 (Figure 3B). Moreover, no ISG induction was observed in PR8 or DIP treated A549 STAT1^-/-^ cells, with the exception of ISG15 and RSAD2 (Viperin), the expression of which was induced by PR8 but not DI-244 (Figure 3B). Finally, results with A549 wt cells were confirmed by qRT-PCR analyses. Induction of IFNβ and IFNλ by DI-244 was detectable but at least 100-fold less efficient as compared to PR8 while differences in ISG induction were frequently less than 10-fold (Figure 3C). In sum, these results suggest that DI-244 inhibits viral infection by STAT1-dependent induction of ISG expression without inducing appreciable expression of IFN.

## Discussion

DI RNAs arise in IAV infected cell cultures, eggs, animals and patients (4, 5, 20, 26-29). They inhibit IAV infection and might modulate IAV intra- and interpatient spread and pathogenesis. However, the mechanism underlying DI RNA antiviral activity and the determinants controlling whether a defective viral genomic RNA is also interfering are incompletely understood. Here, we show that any central deletion in segments 1, 2 and 3 of IAV is sufficient to convert these RNAs into DI RNAs and that inhibitory activity of the respective DI RNAs extends to all tested IAV genomic RNAs. Moreover, we provide evidence that the contribution of replication interference to DIP antiviral activity in cell culture is minor as compared to induction of the IFN system.

IAV and influenza B virus DI RNAs usually contain deletions relative to the genomic RNAs they arose from (4, 5), although an exception has recently been reported (30). Moreover, DI RNAs derived from IAV segments 1-3, which encode the subunits of the viral polymerase, arise more frequently than those derived from other segments (4-6, 8, 9, 31, 32) and were thus in the focus of the present study. The almost universal presence of a deletion in DI RNAs suggests that their shorter length might allow them to out-compete their parental RNAs for resources required for RNA replication. Although this hypothesis is frequently posited (4, 5), direct experimental proof is largely lacking. Here we provide this proof by demonstrating that deleting any internal sequence from segments 1, 2 and 3 is sufficient to generate a DI RNA. Furthermore, we demonstrate that the inhibitory activity of these DI RNAs is determined by their length, at least in the absence of an IFN response, and extends to all target segments tested. The latter observation fits with the finding that DI-244 interferes with replication of several genomic RNAs in IAV infected cells (33). In sum, deleting the sequences between the conserved 5’ and 3’ ends of any IAV RNA, which are required for transcription and translation, will likely generate potent DI RNAs. In some cases, the truncated open reading frame encoded by such DI-RNAs might contribute to antiviral activity (34) but this was not observed for DI-244, in keeping with previous results (33).

Type I IFNs trigger the expression of about 400 genes, many of which encode proteins with antiviral activity, including MX1 (35). The present study shows that when conditions are chosen that allow DIPs to robustly activate the IFN system, DIPs are potent inducers of ISG expression and the contribution of replication interference to DIP antiviral activity is minor. Notably, RNAseq analysis revealed that IAV but not DIPs induced type I and type III IFN expression although both triggered ISG expression in a STAT1-dependent fashion. A potential explanation for this discrepancy is that DIPs induced IFN expression at levels too low to be detected by RNAseq but still sufficient to induce ISGs. Indeed, qRT-PCR analysis revealed modest upregulation of type I and III IFN upon DIP treatment. Alternatively, one might speculate that DIPs may induce ISGs via an unidentified IFN-independent, STAT1-dependent pathway. Interestingly, Wang and colleagues also reported that DIPs induce robust levels of ISGs but only moderate levels of IFN (36) and further research is required to explore why IFN and ISG induction are not correlated in the context of DIPs. Moreover, it is unclear how undiluted DIPs exerted anti-IAV but not anti-VSV activity in STAT1^-/-^ cells without inducing ISGs or other cellular genes. Competition of DIPs with IAV for engagement of entry receptors is one possibility. Collectively, our results underline previous findings that DIPs are potent inducers of antiviral responses (13-17) and show that DIP antiviral activity due to induction of ISG expression outweighs that due to replication interference.

What are the major implications of our findings for DIP development as antivirals and for elucidating the role of naturally occurring DIPs in IAV infection? First, it is essential that antiviral activity of DIPs is examined in IFN competent animal models which express ISGs with potent anti-IAV activity, particularly MX1. Second, antiviral activity due to replication interference can be attained only if DIPs are added in 100 to 1,000 fold excess relative to virus (18) and it remains to be examined whether the strong IFN induction under those conditions exerts unwanted toxic effects in animals and humans. Third, DIP treatment should be more effective in the prophylactic as compared to the therapeutic setting, since only in the former DIP-induced IFN can fully contribute to antiviral activity. Fourth, design of DI RNA and analysis of DI RNAs emerging in patients should focus on the smallest RNAs, since they can be expected to exert the highest antiviral activity.

## Material and Methods

### Plasmids and oligonucleotides

Plasmids for rescue of A/PR/8/34, pHW191-pHW198, and A/WSN/33, pHW181-pHW188, were previously described (37). Plasmids encoding DI RNAs were generated by splice overlap PCR, joining 5’ and 3’-end sequences of desired length, following a strategy previously described for DI-244 (18). A multiple cloning site (mcs) for later insertion of a reporter gene was included in the respective oligonucleotide sequences (Table S2). The PCR products were cloned into pHW2000-GGAarI by golden gate cloning (38). Start codons in DI-244 were mutated using splice overlap PCR primer pairs mutIAV-seg1-ATG-for (5’-TCAATTATATTCAATTTGGAAAGAATAAAAG -3’)/mutIAV-seg1-ATG-rev (5’-CTTTTATTCTTTCCAAATTGAATATAATTGA-3’) and DImut2+3ATG-for (5’-ACTACGAAATCTAATCTCGCAGTCTCGCACCCGCGAGATACTCACAAAAACCACCGTGGACCATATCGCCATAATCAAGAAG-3’)/DImut2+3ATG-rev (5’-CTTCTTGATTATGGCGATATGGTCCACGGTGGTTTTTGTGAGTATCTCGCGGGTGCGAGACTGCGAGATTAGATTTCGTAGT-3’). PCR constructs were cloned into pHW2000-GGAarI as described above.

For expression of the truncated PB2 ORF from DI-244, the ORF was amplified from pHW2000GG-DI244-rep using primers PB2-QCXIP-5N (5’-CCGCGGCCGCACCATGGAAAGAATAAAAGAACTAC-3’)/PB2-3XBgl (5’-GGAGATCTCGAGCTAATTGATGGCCATCCGAAT-3’) digested with NotI/XhoI and cloned into NotI/SalI digested pCAGGS-mcs bearing an altered multiple cloning site (XhoI-SacI-Asp718I-NotI-EcoRV-ClaI-EcoRI-SmaI-SalI-SphI-NheI-BglII).

For generation of empty vector p19polI-GGAarI the insert was amplified from pHW2000-GGAarI by splice overlap PCR using primers HW2-GG-5Bgl, CCdelE-rev (5’-CGTCTTTCATTGCCATACGAAACTCCGGATGAGCATTCATCAG-3’), CCdelE-for (5’-CTGATGAATGCTCATCCGGAGTTTCGTATGGCAATGAAAGACG-3’)/ rRNA-Pr(GG)-3Eco (5’-GCGAATTCTATAGAATAGGGCCAGGTC-3’) and cut with BglII and EcoRI for insertion into BamHI and EcoRI digested p19luc (39).

Reporter plasmids for mini-replicon assay have been described (pPolI-Luc (vRNA/FLUAV/NS1 Seg8-NCR) (19) or were newly generated. First, the reporter with segment 8 ends was amplified with primers fluA AarI-NS-1 and fluA AarI-NS-890R (Table S3) and inserted into vector p19polI-GGAarI by Golden Gate cloning. To generate reporters with ends derived from other segments of IAV, the luciferase reporter gene was amplified with primers encoding the respective untranslated regions (Table S3) and cloned into vector p19polI-GGAarI as described before. All PCR amplified sequences were confirmed by automated sequence analysis.

### Cells and viruses

293T, A549 wt and A549 STAT1^-/-^ cells were maintained in Dulbecco’s Modified Eagle Medium (DMEM; Gibco) containing 10% fetal bovine serum (FBS, Gibco), penicillin (Pen, 100 IU/mL) and streptomycin (Strep, 100 µg/ml). BHK-21 cells were cultivated in Dulbecco’s modified Eagle medium (DMEM, Pan Biotech) supplemented with 5% fetal bovine serum and pen/strep. 293T cell lines stably expressing codon optimized PB2 (293T-PB2opt) were cultured in the presence of 1µg/ml puromycin. Madin-Darby canine kidney cells (MDCK) were cultured in Glasgow’s Modified Eagle Medium (GMEM; Gibco) supplemented with 10% fetal bovine serum (FBS, Gibco) and pen/strep. MDCK cells stably expressing PB2opt were maintained in the presence of 1.5µg/ml puromycin. For generation of A549 STAT1^-/-^ cells, A549 wt cells were transduced with pLentiCRISPR v2 (Addgene, plasmid 52961), a lentivirus expressing Cas9, puromycin resistance, and a guide RNA targeting human *STAT1* (TTCAAGACCAGCGGCCTCTGAGG). Transduced cells were puromycin selected for seven days and surviving cells were plated in 96-well dishes as single cells and expanded. Clonal populations were then lysed and whole cell extract was examined for STAT1 expression by immunoblot. These efforts identified a single clone that demonstrated a complete loss of STAT1 expression, which we refer herein as STAT1^-/-^ cells. All cells lines were regularly tested for mycoplasma contamination.

A/PR/8/34 (H1N1) and A/WSN/33 (H1N1) (37, 40) were produced in embryonated chicken eggs as described previously (41) while A/WSN/33 adapted to growth in A549 cells was obtained from the strain repository of the IVM Münster and was amplified in A549 cells by continuous passaging. IAV titers were determined using focus formation assay as described (18, 38, 42). Replication-competent vesicular stomatitis virus (VSV) expressing eGFP and either wildtype VSV matrix protein (VSV^*^) or a matrix protein variant harboring four amino acid substitutions associated with increased induction of type-I interferon response (VSV^*^M_Q_) have been described elsewhere (43) and were amplified using BHK-21 cells. Further, a VSV glycoprotein trans-complemented, single-cycle VSV replicon that lacks the genetic information for VSV-G but instead codes for eGFP and firefly luciferase genes (VSV^*^ΔG-FLuc) (44) was employed and propagated on BHK-G43 cells (45). All VSV variants were titrated on BHK-21 cells and eGFP-positive foci (replication-competent VSV) or eGFP-positive single cells (single-cycle VSV) were counted as described previously (46).

### Mini-replicon assay

The mini-replicon assay was performed as described (19). In brief, 293T cells seeded in 12-well plates at a cell density of 2 × 10^5^ cells per well were cotransfected with plasmids encoding PB1 (10 ng), PB2 (10 ng), NP (100 ng), reporter segment encoding firefly luciferase (50 ng) and plasmid encoding a DI RNA or empty plasmid (amounts indicated in figures or figure legends). Cells were washed at 6-8 h and harvested at 24 h post transfection. Firefly luciferase activity in cell lysates was measured using a commercial kit (PJK) and the Plate Chameleon V reader (Hidex, Turku, Finland) jointly with Microwin 2000 software.

### Production of DIPs

A coculture of 1.4 ×10^6^ 293T cells and 0.4 ×10^6^ MDCK cells each stably expressing PB2opt and seeded in T-25 flask was cotransfected with plasmids encoding IAV genomic segments 2-8 of either PR8 or WSN origin and a plasmid encoding a segment 1-derived DI-RNA. After overnight incubation, cells were washed once with PBS and, for production of A/PR/8/34-derived DIPs, DMEM infection medium (0.2% MACS BSA, 1% pen/strep) supplemented with TPCK trypsin (0.5 µg/ml) was added. For production of A/WSN/33-derived DIPs, DMEM growth medium (2% FCS, 1% pen/strep) was added. As a negative control, parental MDCK and 293T cells were transfected. Supernatants containing A/PR/8/34-derived DIPs were harvested at 4, 6, 8 and 10 days post transfection while supernatants containing A/WSN/33-derived DIPs were harvested at 3, 5, 7 and 9 days post transfection. Supernatants were cleared from debris by centrifugation, aliquoted and stored at -80 °C for further use. For some experiments, DIPs were further amplified in MDCK-PB2opt cells. For this, a total of 3 ×10^6^ cells were seeded in T-75 flasks and infected at an MOI of 0.01 or lower. Upon detection of CPE, supernatants were cleared from debris by centrifugation and sterile-filtration (0.45 µm filter), aliquoted and stored at -80 °C for further use. Integrity of selected DIP preparations was controlled with segments specific PCR. Infectious titers of supernatants were determined by focus formation assay using MDCK-PB2opt cells as targets, as described (18, 38, 42).

### Analysis of antiviral activity of DIPs

For testing the antiviral activity of DIPs in MDCK cells in the presence of trypsin, cells were seeded at 10,000 cells/well in 96-well plates and coinfected with DIP (MOI 1, and 10-fold dilutions) and IAV (A/PR/8/34, MOI 0.001) for 1 h in Glasgow’s MEM (GMEM) infection medium containing trypsin (0.5 µg/ml). Alternatively, DIPs were added 24 h prior to the virus. For analysis of DIP antiviral activity in MDCK cells, A549 wt and A549 STAT1^-/-^ cells in the absence of trypsin, cells were again seeded at 10,000 cells/well in 96-well plates and either coinfected with DIP (MOI 5 or 10, and 10-fold dilutions) and IAV (A/WSN/33, MOI 0.1) in DMEM medium without trypsin or DIPs added 24 h prior to the virus. After 1 h, cells coexposed to DIPs and virus were washed and culture medium with or without trypsin was added. Supernatants were harvested after 72 h (MDCK) and 96 h (A549 wt and A549 STAT1^-/-^). Viral titers in culture supernatants were quantified using focus formation assay and MDCK cells, as described (38, 42).

### Quantitative RT-PCR analysis

In order to investigate modulation of *MX1* mRNA expression by IAV, DIPs and IFN, a quantitative RT-PCR assay was performed. For this, A549 cells were seeded at a cell density of 2 ×10^5^ cells/well in 12-well plates and inoculated with IAV (MOI 1), DIPs (MOI 1) or pan-IFNα (100 U/ml, PBL Assay Science) using DMEM infection medium for 1 h (DMEM infection medium without trypsin was added to cells exposed to IFNα). Then cells were washed once with PBS and cultured in DMEM infection medium without trypsin for 24 h. To assess the effect of trypsin on *MX1* induction by IFNα, cells were incubated for 24 h with IFNα in the presence of 0, 0.5, 0.05 and 0.005 µg/ml trypsin. At 24 h post treatment, total cellular RNA was extracted using the RNeasy Mini kit (Qiagen) following the manufacturer’s instructions. After determining the RNA content, 1 µg RNA was used as template for cDNA synthesis employing the SuperScript III First-Strand Synthesis System (ThermoFisher Scientific), following the protocol for random hexamers. Subsequently, 1 µl of cDNA (total volume after cDNA synthesis: 20 µl) was analyzed by quantitative PCR on a Rotorgene Q device (Qiagen) employing the QuantiTect SYBR Green PCR Kit (Qiagen). Each sample was analyzed in triplicates for transcript levels - given as cycle threshold (Ct) values - of ß-actin (ACTB, internal transcript control) and myxovirus resistance protein 1 (*MX1*, indicator for IFN induction, target transcript) with primer previously reported by Biesold and colleagues (47). In order to analyze gene expression, the 2-^ΔΔCt^ method was used (48).

The impact of IAV or DIP-on expression of cellular mRNAs was measured by RNAseq and results for certain mRNAs were confirmed by quantitative RT-PCR. Expression of these mRNAs was assayed using TB Green™ Premix Ex Taq™ II (Tli RNase H Plus; Takara) according to manufacturer’s instructions with QuantiTect primer assays (Qiagen). For this, RNA was isolated as described above and reverse transcribed into cDNA with the prime Script RT reagent kit (Takara). 10 ng of cDNA was used in a 1X reaction consisting of 12.5 µl TB Green Premix Ex Taq II (Tli RNaseH plus) (2X), 2 µl 10X QuantiTect primer assay, and 0.5 µl 50X ROX reference dye in a final reaction volume of 25 µl. PCR reactions were performed in a StepOne Plus Instrument (Thermo Fisher). The following QuantiTect primer assays were used. Hs_IFIT1_1_SG (QT00201012), Hs_ISG20_1_SG (QT00225372), Hs_IFNB1_1_SG (QT00203763). Hs_MX1_1_SG (QT00090895), Hs_IFNL1_2_SG (QT01033564), Hs_EIF2AK2_1_SG (QT00022960), Hs_IFNL2_1_SG (QT00222488), Hs_RR18s (QT00199367), Hs_ISG15_1_SG (QT00072814), Hs_RSAD2_1_SG (QT00005271). 18S RNA is used as housekeeping gene. Fold gene induction over mock treated control is calculated by the ΔΔCT method.

### Vesicular Stomatitis Virus Replicon-Based Bioassay

To analyze the relative contribution of IFN induction to antiviral activity, a VSV replicon-based bioassay (44) was performed. This assay is based on the principle that inoculation of effector cells with virus or DIPs leads to the induction of the innate immune system, resulting in the release of type-I IFN into the culture supernatant. These supernatants are then used to inoculate sentinel cells. Here, the type-I IFN will bind to the IFNα/β receptors and trigger a signaling cascade leading to the induction of an antiviral state. Subsequent inoculation of the sentinel cells with a highly IFN-sensitive VSV replicon containing a luciferase reporter will yield luciferase activities that inversely correlate with the extent of the induced antiviral state. A549 and A549 STAT1^-/-^ cells (= effector cells) were seeded in 12-well plates (200,000 cells/well) and inoculated with IAV, VSV^*^, VSV^*^M_Q_, or DIPs (all at MOI of 1) using DMEM infection medium containing trypsin for 1h. The cells were washed once with PBS and cultured in DMEM infection medium without trypsin (used for all further steps) for 16-18 hours. Next, supernatant was harvested and infectious virus was inactivated by addition of 0.1 M HCl and heating the samples for 30 min to 56 °C. After the samples cooled down to room temperature, alkaline treatment was performed using 0.1 M NaOH to neutralize the acidic pH. Subsequently, two-fold serial dilutions of the samples were prepared. In addition, medium containing two-fold serial dilutions of recombinant pan IFNα (starting at a concentration of 400 U/ml) were treated in the same fashion. These samples served as reference and were later used to calculate the relative antiviral activity present in the different supernatants (given as relative IFNα units per ml). The diluted supernatants and IFNα reference samples were added in quadruplicates to a confluent layer of A549 cells grown in 96-well plates (= sentinel cells) and incubated for 24 h. Thereafter, the cells were inoculated with VSV^*^ΔG-FLuc reporter virus (MOI of 3) and further incubated for 6 h. Then, the medium was aspirated and 50 µl/well of 1x luciferase lysis buffer was added. Following an incubation period of 30 min the lysates were transferred into white, opaque-walled 96-well plates and firefly luciferase activity in cell lysates was measured as described above for the mini-replicon assay.

For normalization, luciferase activity was set as 100 % for cells that received regular culture medium instead of diluted culture supernatant/IFNα prior to inoculation with VSV^*^ΔG-FLuc. Using the normalized luciferase values of cells treated with the IFNα reference samples and a non-linear regression model we then calculated the relative IFNα content (given as units per ml) for the effector cell supernatants.

### RNA-seq analysis

For analysis of IAV and DIP mediated modulation of cellular gene expression, A549 wt and A549 STAT1^-/-^ cells were exposed for 1h to DMEM infection medium supplemented with TPCK trypsin (0.5 µg/ml) and containing A/PR/8/34 or DI-244 at an MOI of 1 or were exposed to control supernatants. Subsequently, cells were washed and cultured with DMEM infection medium without trypsin. At 24 h post treatment, total cellular RNA was extracted using the RNeasy Mini kit (Qiagen) following the manufacturer’s instructions and subsequently sent for RNAseq analysis at the Integrative Genomics Core Unit (NIG), Department of Human Genetics, University Medical Center Göttingen.

RNA-seq libraries were performed using the non-stranded mRNA Kit (Illumina). Quality and integrity of RNA was assessed with the Fragment Analyzer using the standard sensitivity RNA Analysis Kit (Advanced Analytical). All samples selected for sequencing exhibited an RNA integrity number of >8. After library generation, we used the QuantiFluor™dsDNA System (Promega) for accurate quantitation of cDNA libraries. The size of final cDNA libraries was determined by using the dsDNA 905 Reagent Kit (Advanced Analytical) exhibiting a sizing of 300 bp in average. Libraries were pooled and sequenced on an Illumina HiSeq 4000 (Illumina) generating 50 bp single-end reads (28-35 Mio reads/sample). The raw read & quality check were done by transforming sequence images the BaseCaller software (Illumina) to BCL files, whichwere demultiplexed to fastq files with bcl2fastq v2.20. The sequencing quality was asserted using FastQC (http://www.bioinformatics.babraham.ac.uk/projects/fastqc/).

For subsequent data analysis, ISGs with anti-IAV activity were selected based on work by Schoggins and colleagues (35). ISG expression in IAV- or DIP-treated cells was further normalized to ISG expression in control-treated cells.

## Acknowledgements

We would like to thank Benjamin tenOever and Martin Schwemmle for the kind gift of A549 STAT1^-/-^ cells and minireplicon plasmids, respectively. This study was supported by the following grants: Defense Advanced Research Projects Agency (DARPA) to SP and UR, Bundesministerium für Bildung und Forschung (BMBF, grant number 01KI1723E) to SP and the European Union’s Horizon 2020 research and innovation programme under grant agreement number 101003666 (OPENCORONA) to FW.

